# Photoperiod stress alters the cellular redox status and is associated with an increased peroxidase and decreased catalase activity

**DOI:** 10.1101/2020.03.05.978270

**Authors:** Walid Abuelsoud, Anne Cortleven, Thomas Schmülling

**Author notes:** **E-mail addresses**: WA,; AC,; TS. **Corresponding authors**: Prof. Dr. Thomas Schmülling and Dr. Anne Cortleven, Institute of Biology/Applied Genetics, Dahlem Centre of Plant Sciences (DCPS), Freie Universität Berlin, Albrecht-Thaer-Weg 6, D-14195 Berlin, Germany, / /+49 30 838 56796.

## Abstract

Periodic changes of light and dark regulate numerous processes in plants. Recently, a novel type of stress caused by an extended light period has been discovered in *Arabidopsis* and was named photoperiod stress. Photoperiod stress causes the induction of numerous stress response genes during the night following the extended light period of which many are indicators of oxidative stress. The next day, stress-sensitive genotypes display reduced photosynthetic efficiency and programmed cell death in leaves. Here, we have analysed further the consequences of photoperiod stress and report that it causes changes of the cellular redox status. A prolonged light period caused a strong reduction of the AsA redox during the following night indicating that it induces an oxidizing cellular environment. Further, photoperiod stress was associated with an increased activity of peroxidases and a decreased activity of catalases. Increased peroxidase activity was localized to the apoplast and might be causal for the oxidative stress induced by photoperiod stress.

## Introduction

Plants are exposed to a regular daily light/dark rhythm. Changes in this rhythm due to changes in the photoperiod have a strong impact on many biochemical, physiological and developmental processes during plant life including flowering and hypocotyl growth, as well as abiotic and biotic stress responses (Greenham and McClung, 2015; Shim and Imaizumi, 2015).

Recently it has been described that changes of the photoperiod, in particular a prolongation of the light period, induce a stress response during the following night. This new form of abiotic stress was named photoperiod stress (originally circadian stress) (Nitschke *et al.*, 2016; Nitschke *et al.*, 2017). The stress phenotype was discovered in plants with a reduced cytokinin (CK) content or signaling as these showed a particularly strong photoperiod stress response. This response occurs during the night following an extended light period and includes a strong induction of stress marker gene expression and increase of jasmonic acid content. The following day, a reduced photosynthetic efficiency and eventually programmed cell death (PCD) in leaves ensues. Induction of stress marker genes indicated that wild-type plants also receive and respond to this stress but they showed only a weak or no leaf phenotype. It was concluded that CK is required to protect against photoperiod stress (Nitschke *et al.*, 2016).

CKs are known to play a central role in many physiological and developmental processes during plant life (Werner and Schmülling, 2009; Kieber and Schaller, 2018) and in the response to biotic and abiotic stresses (Cortleven *et al.*, 2019). In *Arabidopsis*, the CK signal is perceived by three different receptors, namely ARABIDOPSIS HISTIDINE KINASE2 (AHK2), AHK3, and CYTOKININ RESPONSE1 (CRE1)/AHK4 (Inoue *et al.*, 2001; Suzuki *et al.*, 2001). Upon CK perception, signal transduction takes place through a multistep phosphorelay mechanism similar to the bacterial two-component system to regulate the expression of CK response genes (Heyl *et al.*, 2012). In the response to photoperiod stress CK acts through the receptor AHK3 and the B-type response regulators ARR2, ARR10 and ARR12 (Nitschke *et al.*, 2016).

Besides CK-deficient plants, also certain clock mutants showed a strong response to photoperiod stress. Common to stress-sensitive clock mutants and CK-deficient plants was a lowered expression or impaired functioning of CIRCADIAN CLOCK ASSOCIATED1 (CCA1) and LONG HYPOCOTYL (LHY), two key components of the morning loop (for review see Shim and Imaizumi, 2015), which indicated that a functional clock is also essential to cope with stress caused by altered light-dark rhythms (Nitschke *et al.*, 2016). The clock is necessary to achieve synchronization of internal diurnal processes with the environment. Clock output genes control, together with environmental cues like light, numerous physiological and developmental processes such as flowering time, growth, stomatal movement, redox homeostasis as well as the response to biotic and abiotic stresses (Greenham and McClung, 2015; Karapetyan and Dong, 2018).

Nitschke et al. (2016) showed that photoperiod stress causes oxidative stress as indicated by lipid peroxidation and stress symptoms typically associated with the formation of reactive oxygen species (ROS). ROS, which include hydrogen peroxide (H_2_O_2_), superoxide, hydroxyl radicals and singlet oxygen, are unavoidable toxic versatile byproducts of aerobic metabolism (Mignolet-Spruyt *et al.*, 2016). They are well known for their roles in abiotic and biotic stress responses (Foyer and Noctor, 2013; Xiong *et al.*, 2015; Schmidt *et al.*, 2016; Mhamdi and Van Breusegem, 2018). ROS are highly reactive and can damage many cellular compounds. Therefore, plants have developed various enzymatic and non-enzymatic ROS scavenging systems to maintain ROS homeostasis and manage oxidative stress (Asada, 2006; Sharma and Duda, 2012). More recently, ROS have also been recognized as important regulators of growth and development and a distinction has been made between toxic and beneficial levels of ROS (Mittler, 2017; Noctor *et al.*, 2018). ROS signaling is controlled by a highly regulated ROS production in different cellular compartments including mitochondria, chloroplast, peroxisomes and the apoplast (Noctor and Foyer, 2016). ROS production in the apoplast occurs through the membrane-bound RESPIRATORY BURST OXIDASE HOMOLOGS (RBOH) family, which are NADPH oxidases (Dubiella *et al.*, 2013), and apoplastic peroxidases (PRX) (O’Brien *et al.*, 2012; Qi *et al.*, 2017). The *Arabidopsis* genome encodes 73 class III peroxidases, of which the great majority has been predicted to be localized to the apoplast (Valerio *et al.*, 2004). Some of these - PRX4, PRX33, PRX34 and PRX71 - are involved in mediating stomatal resistance against bacteria in a CK-mediated manner (Arnaud *et al.*, 2017).

The initial study on photoperiod stress by Nitschke *et al.* (2016) did not show an increase in H_2_O_2_ concentration as measured by amplex red indicating that probably ROS other than H_2_O_2_ were responsible for the oxidative stress response. In order to learn more about the impact of photoperiod stress on the cellular redox system we have analyzed in more detail the changes in redox status and the enzymatic and non-enzymatic scavenging mechanisms after photoperiod stress. Our results revealed that the oxidative stress resulting from photoperiod stress reduces the AsA redox and is associated with a reduced activity of catalase (CAT) and an enhanced activity of apoplastic PRX, which is unusual for a response to abiotic stress.

## Material and methods

### Plant material and growth conditions

*Arabidopsis thaliana* accession Col-0 was used as wild type (WT). The CK receptor mutant *ahk2-5 ahk3-7* (Riefler *et al.*, 2006) and the clock mutant *cca1-1 lhy-20* (Nitschke *et al.*, 2016) were described before. *Arabidopsis* plants were grown on soil under short day (SD) conditions (8 h light/16h dark) in a growth chamber with light intensities of 100 to 150 μmol m^−2^ s^−1^, using a combination of Philips Son-T Agros 400W, and Philips Master HPI-T Plus, 400W/645 lamps, at 22°C and 60% relative humidity.

### Stress treatment

For stress treatments, five-week-old SD-grown plants were used. The standard stress regime consisted of a 32 h light treatment (prolonged light, PL) integrated into a SD regime (Fig. 1A). Control plants remained under SD conditions. For phenotypical analyses, leaves from stress-treated plants of the same developmental stage were chosen. For RNA and biochemical measurements, only the distal halves of these leaves (leaves 6 - 10) were harvested, flash frozen and homogenized with a Retsch Mixer Mill MM2000 (Retsch, Haan, Germany) with two stainless-steel beads (2 mm diameter). Whole leaves were used for electrolyte leakage (EL) and Fv/Fm measurements. Harvest during the dark period was performed in green light.

**Fig. 1.**
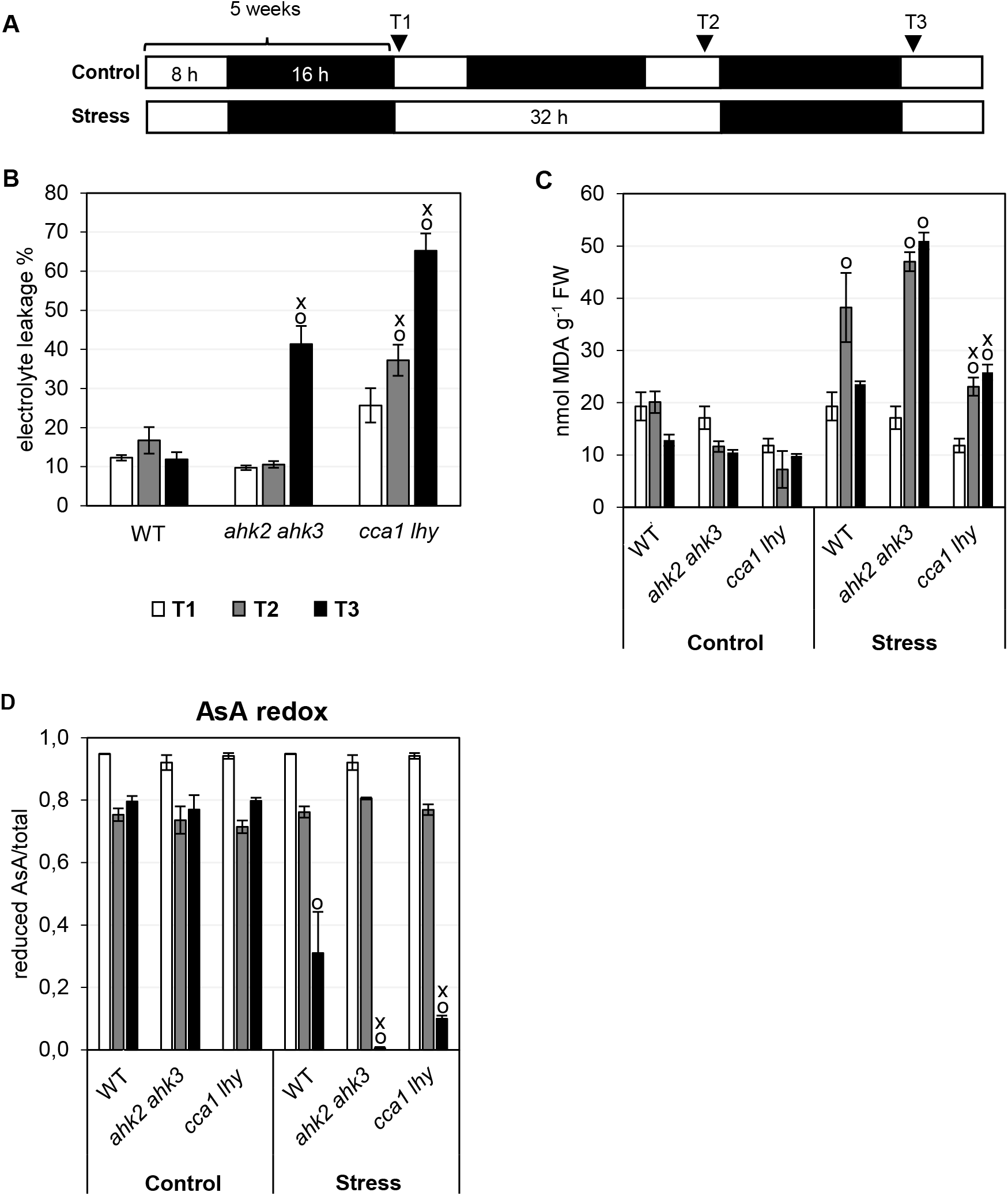
Photoperiod stress is associated with oxidative stress. (A) Schematic overview of the experimental setup used in (B - D). Plants were grown under short day conditions for five weeks and then exposed to a 32-hours light period. Leaf samples were collected at the indicated time points (triangles). White, light period; black, dark period. (B) electrolyte leakage (n = 10), (C) MDA levels (n = 4), (D) Ascorbic acid (AsA) redox (n =4) in leaves at time points indicated in (A). Data are mean values ± SE. Symbols indicate significant differences from the corresponding control (o) and the respective wild type under the same condition (x) (p < 0.05; *t-* test). FW, fresh weight.

### Analysis of cell death

Mature leaves, defined as fully expanded leaves, with or without lesions were counted 20 to 24 h after PL treatment. Percentage of lesions means the percentage of mature leaves with lesions.

### Analysis of photosynthetic efficiency

Chlorophyll fluorescence emission was measured on detached leaves with a modulated chlorophyll fluorometer (Photosystem Instruments, Drasov, Czech Republic). After dark adaptation for 20 minutes, the maximal photochemical efficiency of PSII was determined from the ratio of variable (F_V_) to maximum (F_M_) fluorescence [F_V_/F_M_ = (F_M_-F_0_)/F_M_]. An actinic light pulse (0.2 μmol m^−2^ s^−1^) was used to determine the initial (minimum) PSII fluorescence in the dark-adapted state (F_0_), and F_M_ was determined by a saturating light pulse (1.500 μmol mol^−2^ s^−1^).

### Electrolyte leakage

Membrane leakage of leaves was measured according to Lutts *et al.* (1995). Whole leaves were gently washed to remove any solutes from surfaces, incubated in 20 ml of deionized water at room temperature for 18 h while gently shaking and then boiled in a water bath for 30 min. The conductivity of the solution was measured with a conductivity meter and relative electrolyte leakage (EL) calculated as percentage of initial to final conductivity.

### Malondialdehyde (MDA)

MDA levels were measured according to Hodges (1999). Briefly, 500 μl 0.1% cold TCA was added to the harvested leaf material. After centrifugation at 10.000 *g* for 15 min at 4 °C, the supernatant was incubated with thiobarbituric acid (TBA), to produce thiobarbituric acid-malondialdehyde (TBA-MDA). Absorbance was measured at 440, 532 and 600 nm in a 96-well plate reader (Synergy HT, Biotek, Vermont, USA).

### Total phenolics and flavonoids

Polyphenols and flavonoids were extracted from leaf material in 1 ml 80% methanol (v/v) during centrifugation at 10.000 *g* for 15 min at 4 °C. Total phenolic content was determined using a Folin-Ciocalteu assay according to Zhang *et al.* (2006) and adapted to a 96-well microplate as described in Boestfleisch *et al.* (2014). Gallic acid was used as a standard. Flavonoid content was estimated using the modified aluminum chloride colorimetric method and adapted to a 96-well microplate as described in Chang *et al.* (2002) and Boestfleisch *et al.* (2014) with quercetin as standard.

### Total antioxidant capacity (TAC)

100 mg of fresh finely ground leaf tissues was extracted by the addition of 1 ml ice-cold 80% (w/w) ethanol. TAC of the extract was measured by using FRAP (ferric reducing antioxidant power) reagent according to Benzie and Strain (1999).

### Extraction and assay of ascorbate

Leaf material was extracted in 500 μl 5% TCA and after centrifugation the supernatant was used for assaying the reduced and total ascorbate content according to Boestfleisch *et al.* (2014).

### Extraction and assay of antioxidant enzymes

Activities of APX (EC 1.11.1.11), DHAR (EC 1.8.5.1), MDHAR (EC1.6.5.4), GR (EC 1.8.1.7), SOD (EC 1.15.1.1), catalase (EC1.11.1.6), NADPH oxidase (EC 1.6.3.1) and PRX (EC 1.11.1) were measured in leaf material extracted with 1 mL of ice-cold 50 mM MES-KOH buffer (pH 6.0) containing 40 mM KCl and 2 mM CaCl_2_ followed by vortexing and centrifugation at 16.000 *g* for 20 min at 4°C. 1 mM L-ascorbic acid was added to the extraction buffer when ascorbate peroxidase was extracted. All enzyme assays were performed in a final volume of 0.2 mL in a 96-well microplate at 25 °C (PowerWave HT microplate spectrophotometer; BioTek, Vermont, USA). Samples and blanks were analyzed in triplicate. SOD activity was determined according to Dhindsa *et al.* (1981) by measuring the inhibition of NBT (nitroblue tetrazolium) reduction at 560 nm. 50% inhibition was considered as 1 unit of enzyme. PRX activity was determined by monitoring the oxidation of guaiacol (ε_470_ = 26.6 mM^−1^ cm^−1^) in 50 mM K-phosphate pH 6.0 containing 25 mM H_2_O_2_ and 25 mM guaiacol (Kumar and Khan, 1982). CAT activity was assayed by monitoring the decomposition of H_2_O_2_ (ε_240_ = 43.6 M^−1^ cm^−1^) at 240 nm in 50 mM K-phosphate buffer pH 7.0 containing 25 mM H_2_O_2_ (Aebi, 1984). APX, MDHAR, DHAR and GR activities were measured by the methods of Murshed *et al.* (2008). APX activity was estimated by following the change in absorbance at 290 nm due to oxidation of AsA in a reaction mixture containing 50 mM K-phosphate buffer pH 7.0, 0.25 mM AsA and 5 mM H_2_O_2_ (ε_ascorbic acid_ = 2.8 mM^−1^ cm^−1^). The DHAR reaction is started by the addition of freshly prepared DHA to a final concentration of 0.2 mM in 50 mM HEPES buffer (pH 7.0) into all wells and following the increase in absorbance at 265 nm for 5 min. Specific activity was calculated from the 14 mM^−1^ cm^−1^ extinction coefficient. MDHAR activity was assayed in 50 mM HEPES buffer pH 7.6 containing 2.5 mM AsA, 0.25 mM NADH and 0.4 U of ascorbate oxidase. The activity is measured by following the decrease in absorbance at 340 nm (ε = 6.22 mM^−1^ cm^−1^). GR was assayed in a reaction mixture containing 50 mM HEPES buffer pH 8.0 and containing 0.5 mM GSSH, 0.5 mM EDTA, 0.25 mM NADPH. The activity was calculated by monitoring decrease in absorbance at 340 nm and by using the extinction coefficient 6.22 mM^−1^ cm^−1^.

### Apoplastic peroxidase activity

Extraction of the apoplastic solution from leaf material was carried out according to Córdoba-Pedregosa *et al.* (2004) and detailed in Araya *et al.* (2015). Distal halves of 12 leaves (for WT and *cca1 lhy*) and 16 leaves (for *ahk2 ahk3*) were harvested, quickly washed in distilled water, the surface was gently wiped with soft paper towels and placed in Petri dishes submerged in 10 mM sodium phosphate buffer, pH 6, containing 1.5% polyvinylpolypyrrolidone, 1 mM EDTA and 0.5 mM phenylmethylsulphonyl fluoride, and then submitted to vacuum (60 kPa) for 5 min at 4 °C. Then, the surface of leaves was dried with soft paper towels and placed in syringes, which were then placed in falcon tubes. After 150 *g* centrifugation for 5 min, the apoplastic fluids recovered at the bottom of the tubes. Cytosolic contamination of apoplastic solution was monitored by assaying glucose-6-phosphate dehydrogenase (G6PDH) activity as a marker of cytoplasmic contamination according to Córdoba-Pedregosa *et al.* (2004). PRX activities were assayed as described above.

### Cell wall-bound peroxidase activity

The cell wall fraction was extracted from leaf material by addition of ice-cold 50 mM phosphate buffer (pH 5.8) followed by centrifugation for 15 min at 10.000 *g* at 4°C and four times washing of the pellet with extraction buffer as described in Lin and Kao (2001). PRX which is ionically bound to cell walls was extracted by incubating the cell wall preparation in ice-cold 1 M NaCl in 50 mM phosphate buffer (pH 7) for 2 h while shaking and assayed as described above.

### Determination of protein concentrations

The protein content in the enzyme extracts was determined by using Bradford assay (BioRad) (Bradford, 1976).

### RNA isolation and quantitative real-time RT-PCR

Total RNA was extracted from leaf material using the NucleoSpin^®^ RNA plant kit (Machery and Nagel, Düren, Germany) as described in the user’s manual or by a phenol/chloroform/LiCl isolation adapted from Sambrook and Russell (2001). Shortly, RNA was extracted from frozen leaf material by the addition of 750 μl extraction buffer (0.6 M NaCl, 10 mM EDTA, 4 % (w/v) SDS, 100 mM Tris/HCl pH 8) and 750 μL phenol/chloroform/isoamyl alcohol (25:24:1). Samples were vortexed, shaken for 10 min at RT and centrifuged at 19.000 *g* for 5 min at 4 °C. The supernatant was transferred into a fresh 1.5 mL Eppendorf tube and chloroform/isoamyl alcohol (24:1) was added in a 1:1 ratio. After centrifugation at 19.000 *g* for 5 min at 4 °C, the RNA was precipitated for 2 h on ice by adding 0.75 volumes of 8 M LiCl. After centrifugation at 19.000 *g* for 15 minutes at 4 °C, supernatant was resolved in 300 μL RNase-free water and RNA was precipitated again by the addition of 30 μL 3 M sodium acetate and 750 μL absolute ethanol during incubation at −70 °C for 30 min. After centrifugation, the pellet was washed with 200 μL 70% ethanol, dried and resolved in 40 μL RNase-free water.

The RNA concentration was determined spectrophotometrically at 260 nm using a Nanodrop ND-1000 spectrophotometer (Nanodrop Technologies, Wilmington, USA). The RNA purity was evaluated by measuring the 260/280 nm ratio. After a DNAse treatment (Fermentas, Life Technologies, Darmstadt, Germany), equal amounts of starting material (1 μg RNA) were used in a 20 μl SuperScript® III Reverse Transcriptase reaction. First strand cDNA synthesis was primed with a combination of oligo(dT)-primers and random hexamers. Primer pairs were designed using Primer 3 Software (http://www.genome.wi.mit.edu/cgibin/primer/primer3.cgi) or Quantprime Software (Arvidsson *et al.*, 2008) under the following conditions: optimum Tm at 60 °C, GC content between 20% and 80%, 150 bp maximum length. Primers used are listed in Table S1. Quantitative real-time RT-PCR using FAST SYBR Green I technology was performed on an CFX96 Touch Real-Time Detection System (Biorad, Feldkirchen, Germany) using standard cycling conditions (15 min 95°C, 40 cycles of 5 s at 95°C, and 15 s at 55°C and 10 s at 72°C) followed by the generation of a dissociation curve to check for specificity of the amplification. Reactions contained SYBR Green, Immolase (Bioline, Memphis, USA), 300 nM of a gene-specific forward and reverse primer and 2 μl of a 1:10 diluted cDNA in a 20 μl reaction. Gene expression data were normalized against two or three different nuclear-encoded reference genes (*UBC21, PP2A* and/or *MCP2A*) according to Vandesompele *et al.* (2002) and presented relative to the level in WT at time point 1.

### Statistical analysis

Statistical analyses were conducted using GraphPad Prism 8 statistical software. Significant differences between the means were tested by Student’s *t*-test at *P* ≤ 0.05

## Results

### Photoperiod stress induces oxidative stress

To induce photoperiod stress, we used the standard stress regime from Nitschke *et al.* (2016) which consists of a 32 h prolonged light (PL) period given to five-week-old short day-adapted plants (Fig. 1A). This treatment caused a very strong stress syndrome in the particularly sensitive CK-deficient plants and certain clock mutants. In this study, the stress response of the CK receptor mutant *ahk2 ahk3* and the clock mutant *cca1 lhy* were analyzed in more detail and compared to the much weaker response of WT plants.

In a first approach, we explored the impact of photoperiod stress on the redox status and took samples at the beginning and end of the extended light period and after the end of the following night (Fig. 1A). The stress treatment resulted in a strong increase in lesion formation and a strong decrease in photosynthetic capacity (Fv/Fm) in both *ahk2 ahk3* and *cca1 lhy* mutants (Supplemental Fig. S1A-B). A strong increase in electrolyte leakage was observed in the stress-sensitive *ahk2 ahk3* and *cca1 lhy* mutants the day following photoperiod stress treatment (Fig. 1B). The levels of malondialdehyde (MDA), which is an indicator of lipid peroxidation, were increased at the end of the light treatment both in WT and in the mutants. However, thereafter they decreased in WT but remained high in both *ahk2 ahk3* and *cca1 lhy* mutants (Fig. 1C). These observations are in accordance with the results described in Nitschke *et al.* (2016) and are indicative of oxidative stress.

In order to study the cellular redox state, the AsA (ascorbic acid) redox (ratio of reduced form to total amount) was determined (Fig.1, Suppl. Table S2). AsA redox was not changed at the end of the PL period but decreased strongly in all genotypes during the following night, but much stronger in *ahk2 ahk3* and *cca1 lhy* mutants than in WT (Fig. 1D). This results indicates an oxidizing cellular environment in the mutants’ tissues in response to photoperiod stress.

Furthermore, non-enzymatic antioxidants and activities of scavenging enzymes were measured together with the total antioxidant capacity (FRAP). Both the FRAP and phenolics content showed a small increase in WT and mutant plants after the PL period which increased even further after the following night (Fig. 2A, B). This increase was slightly higher in *cca1 lhy*. The flavonoid content was not strongly altered in any of the genotypes (Fig. 2C). These results point to a rather minor role of non-enzymatic antioxidants to protect plants from oxidative stress caused by photoperiod stress.

**Fig. 2.**
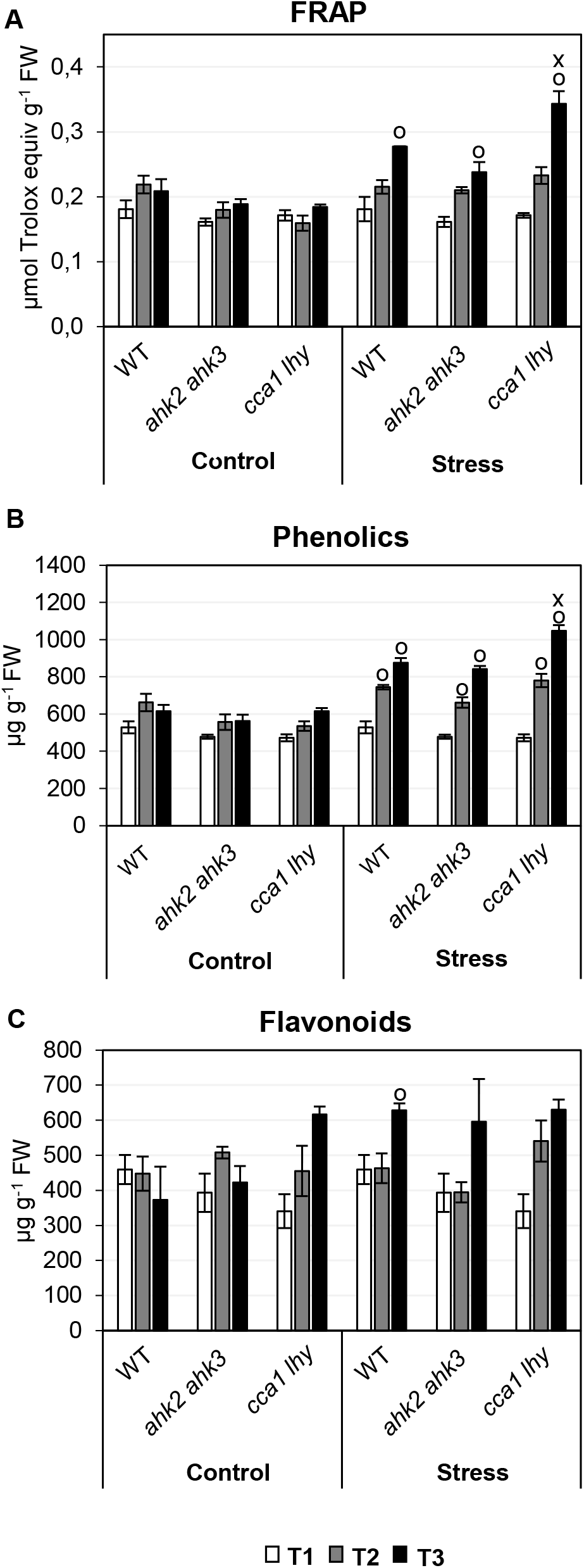
Changes in non-enzymatic antioxidants in response to photoperiod stress. Total antioxidant capacity (A), phenolics (B) and flavonoids (C) in leaves of WT, *ahk2 ahk3* and *cca1 lhy* plants under control and photoperiod stress conditions. Experimental design is described in Fig. 1A. Data are mean values ± SE (n = 4). Symbols indicate significant differences from the corresponding control (o) and the respective wild type under the same condition (x) (p-values < 0.05; *t-*test). FW, fresh weight.

Among the enzymatic antioxidants, APX, MDHAR, DHAR, GR and SOD showed only slight or no significant differences in *ahk2 ahk3* and *cca1 lhy* plants compared to WT both before and after stress treatment (Fig. 3A–E) indicating that these scavenging enzymes are not relevant for the photoperiod stress response. In contrast, the enzyme activities of both CAT and PRX changed strongly in response to photoperiod stress in *ahk2 ahk3* and *cca1 lhy* mutants (Fig. 3F, G). CAT activity strongly decreased to about 20% of its original level while PRX activity significantly increased more than two-fold in *ahk2 ahk3* and *cca1 lhy20* leaves after dark relaxation.

**Fig. 3.**
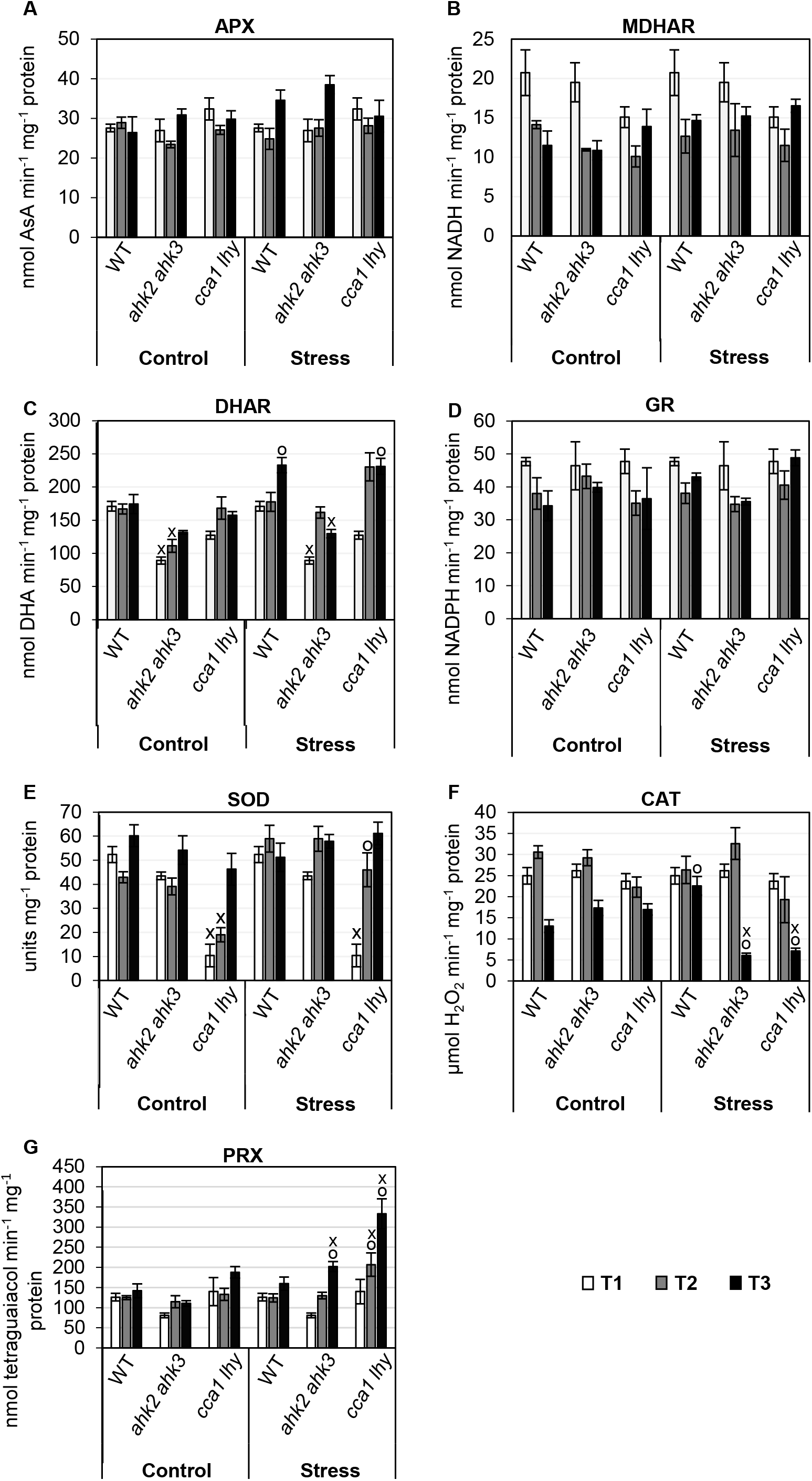
Changes in enzymatic antioxidant activity in leaves of WT, *ahk2 ahk3* and *cca1 lhy* plants in response to photoperiod stress. (A) ascorbate peroxidase (APX), (B) monodehydroascorbate dehydrogenase (MDHAR), (C) dehydroascorbate reductase (DHAR), (D) glutathione reductase (GR), (E) superoxide peroxidase (SOD), (F) catalase (CAT) and (G) peroxidase (PRX) activities under control and stress conditions. Experimental design is described in Fig. 1A. Data are mean values ± SE, n = 4. Symbols indicate significant differences from the corresponding control (o) or the respective wild type under the same conditions and time point (x) (p-values < 0.05; *t-*test).

Together, these results indicated that *ahk2 ahk3* and *cca1 lhy* and to a lesser extent also WT experience oxidative stress as a consequence of photoperiod stress. This oxidative stress occurred during the night following the PL treatment. It was associated with decreased CAT and increased PRX activities which might be causative for the stress. Next we studied the development of the oxidative stress during the night following the extended dark period in a more detailed time course.

### Activities of catalase and peroxidase change during dark relaxation

To investigate at which time point during dark relaxation the oxidative stress starts, we collected samples from the leaves before, during and after dark relaxation (Fig. 4A) and determined the activities of the scavenging enzymes at different time points during the dark. The activities of SOD, APX, DHAR, MDHAR and GR were not different in plants treated by photoperiod stress as compared to control plants (Supplemental Fig. S2).

**Fig. 4.**
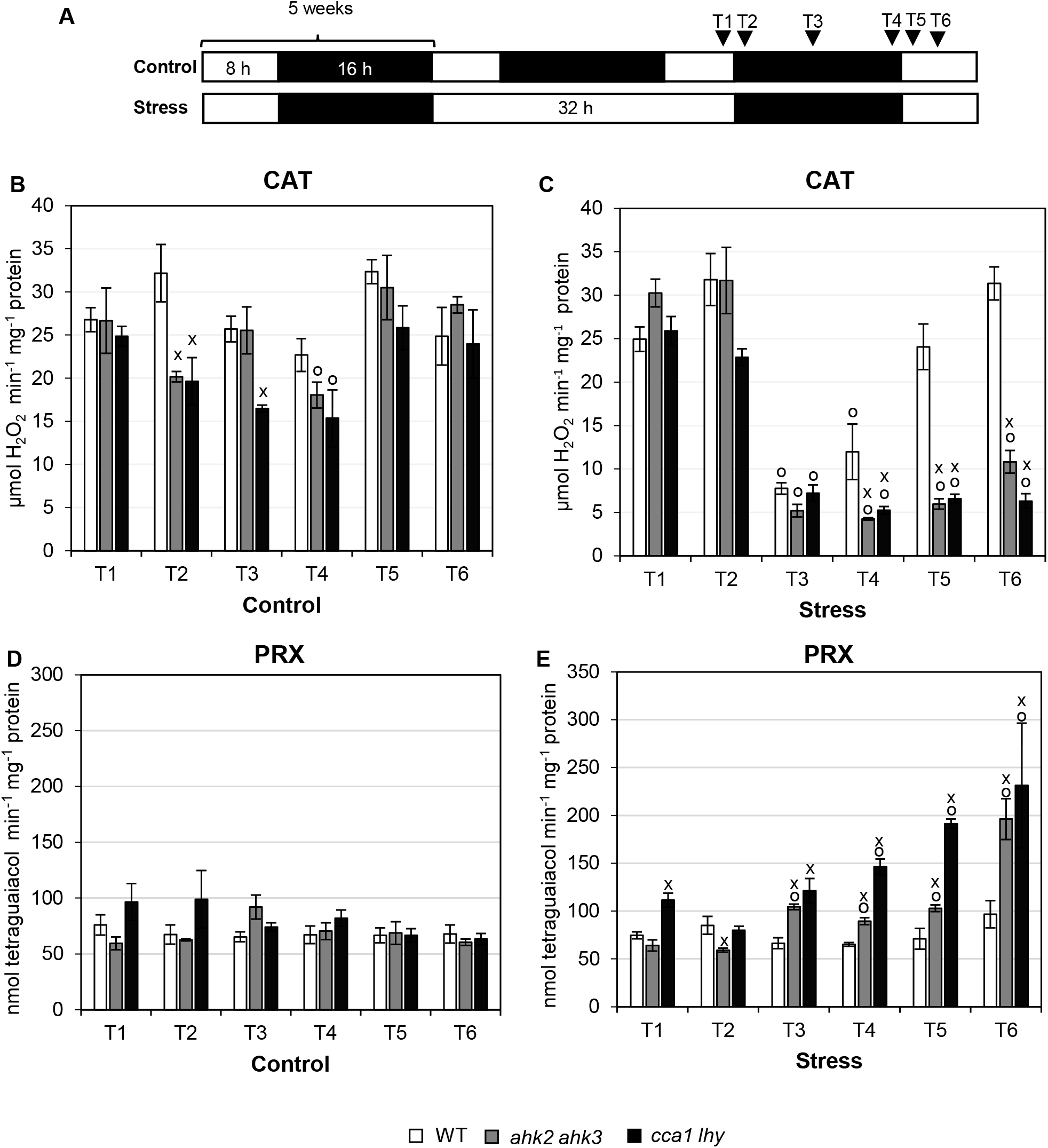
Changes in catalase and peroxidase activities in response to photoperiod stress. (A) Schematic overview of the experimental setup used in (B-E). Catalase (CAT) (B, C) and peroxidase (PRX) (D, E) activity in leaves of WT, *ahk2 ahk3* and *cca1 lhy* plants under control conditions (B, D) and in response to photoperiod stress (C, E) at time points indicated in Fig. 4A. Data are mean values (n = 4; ± SE). Symbols indicate significant differences from the corresponding plants at time point T1 (o) or the respective wild type under the same condition and time point (x) (p-values < 0.05; *t-*test).

The results reported above indicated that especially CAT and PRX might play an important role in the onset of the oxidative stress response. The time course of CAT and PRX activities showed distinct changes in response to photoperiod stress (Fig. 4). CAT activity showed a remarkable reduction in all PL-treated plants after 8 h of dark relaxation (T3). In WT plants CAT activity gradually returned to its original value during and after dark relaxation, while it remained at a low level in the mutant plants (Fig. 4B, C). PRX activity, on the other hand, started to increase in stress-treated *ahk2 ahk3* and *cca1 lhy* leaves after 8 h of dark relaxation and continued to increase further during the night and even during the following light period. No significant changes in PRX activity were noted in control plants or stress-treated WT plants (Fig. 5D, E). Together, these results strongly suggest that the oxidative stress might be caused at least partially by the decreased catalase and increased peroxidase activity.

**Fig. 5.**
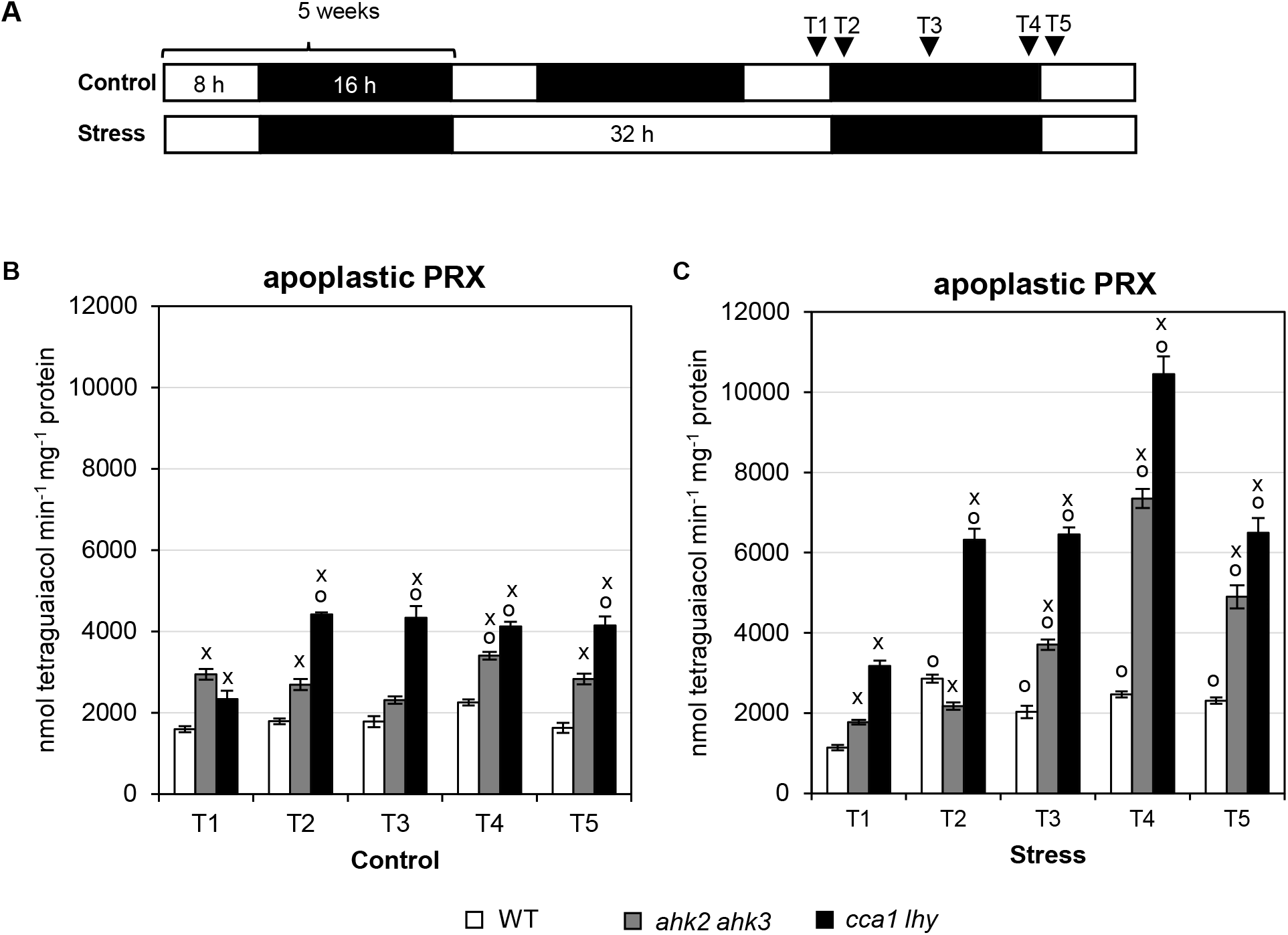
Changes in apoplastic peroxidase activity in response to photoperiod stress. (A) Schematic overview of the experimental setup. Plants (WT, *ahk2 ahk3*, *cca1 lhy*) were grown under short day conditions for five weeks and then exposed to a 32-hours light period. Leaf samples were collected at the indicated time points (triangles). White, light period; black, dark period. (B, C) Apoplastic peroxidase (PRX) activity in leaves of plants grown under control conditions (B) and in response to photoperiod stress (C). Data are mean values ± SE (n = 4). Symbols indicate significant differences from the corresponding plants at time point T1 (o) or the respective wild type under the same condition and time point (x) (p-values < 0.05; *t-*test).

To test if the changes in PRX activity were eventually caused by alteration of apoplastic PRX, the apoplastic solution was extracted from leaves at different time points during dark relaxation (Fig. 5A) and PRX activity were measured. To ensure the purity of the apoplastic extraction, glucose-6-phosphate dehydrogenase (G6P) activity was assayed. The results showed almost no activity of G6P indicating a pure apoplastic fluid (Suppl. Table S3). Apoplastic PRX activity did not change a lot at different time points under control conditions (Fig. 5). Upon photoperiod stress, the apoplastic PRX activity was increased by about twofold in WT and about four- and fivefold in CK receptor and clock mutants (Fig. 5). The cell wall-associated PRXs do not seem to contribute to the oxidative burst since their activity did not change during dark relaxation (Supplemental Fig. S3).

In addition to enzymatic activities, also the transcript levels of *CAT* and *PRX* genes were analyzed at different time points during dark relaxation (Fig. 6A). Our data show that transcript levels of *CAT1* and *CAT3* behave similar: These were high at the beginning of the dark period and decreased gradually over time in both control and stress-treated plants (Fig. 6B,D). In the clock mutant, transcript levels were strongly reduced in comparison to WT and the *ahk2 ahk3* mutant. In contrast, *CAT2* transcripts levels were low at the beginning of the night and showed a gradual increase under control conditions for all genotypes. After stress treatment, this gradual increase was completely missing in all genotypes, they were even further decreased in the mutant plants (Fig. 6C).

**Fig. 6.**
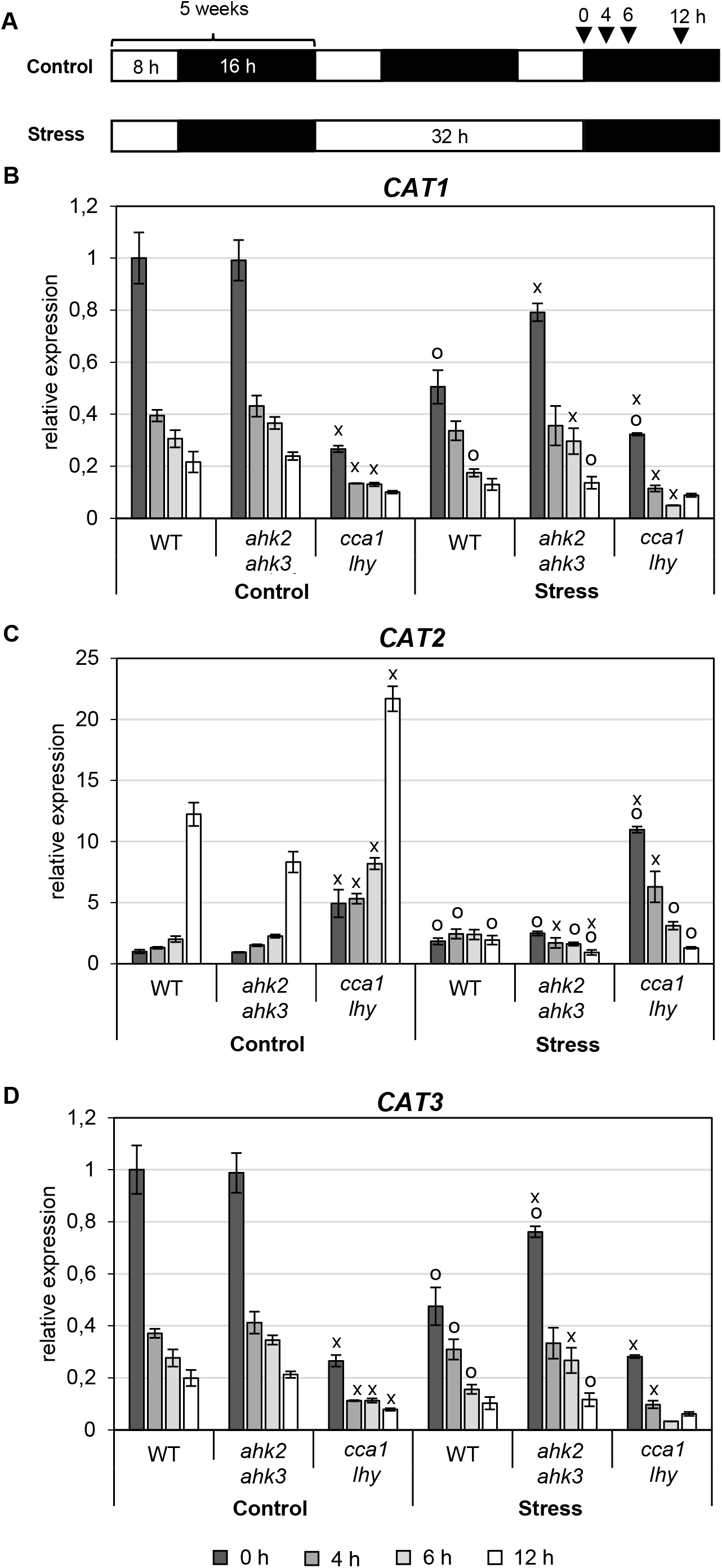
Expression of catalase genes in response to photoperiod stress. (A) Schematic overview of the experimental setup used in (B-D). Plants were grown under short day conditions for five weeks and then exposed to a 32-hours light period. Leaf samples were collected at the indicated time points (triangles). White, light period; black, dark period. (B-D) Transcript abundances of *CATALASE1* (*CAT1*) (B), *CAT2* (C) and *CAT3* (D) in leaves at the time points indicated in (A). Transcript levels were normalized to the 0 h wild-type control, which was set to 1. Data are mean values ± SE (n = 4). Symbols indicate significant differences from the corresponding control (o) or the respective wild type under the same condition and time point (x) (p-values < 0.05; *t-*test).

Also, the transcript levels of *PRX* genes (*PRX4*, *PRX33*, *PRX34, PRX71*) were analyzed. Selection of these genes was based on Arnaud *et al.* (2017) who showed a connection between CK, these *PRX* genes and ROS production. All four genes showed a response to photoperiod stress. Under control conditions, steady state mRNA levels were generally low and decreased slightly during the night. Upon photoperiod stress treatment *PRX* gene expression increased gradually during the night although with different kinetics. *PRX4* responded the fastest and started to increase 4 h after beginning of the night (particularly strong in *cca1 lhy* with already a 400-fold increase at that time), its expression peaked at 6 h and then declined rapidly (Fig. 7A). *PRX71* also responded fast but the induction level was much lower than for *PRX4* (Fig. 7D). *PRX33* and *PRX34* levels increased only later reaching a 4-5-fold increase 12 h after onset of darkness (Fig. 7B, C). Noteworthy, no major differences in transcript levels were observed between the genotypes for these two genes.

**Fig. 7.**
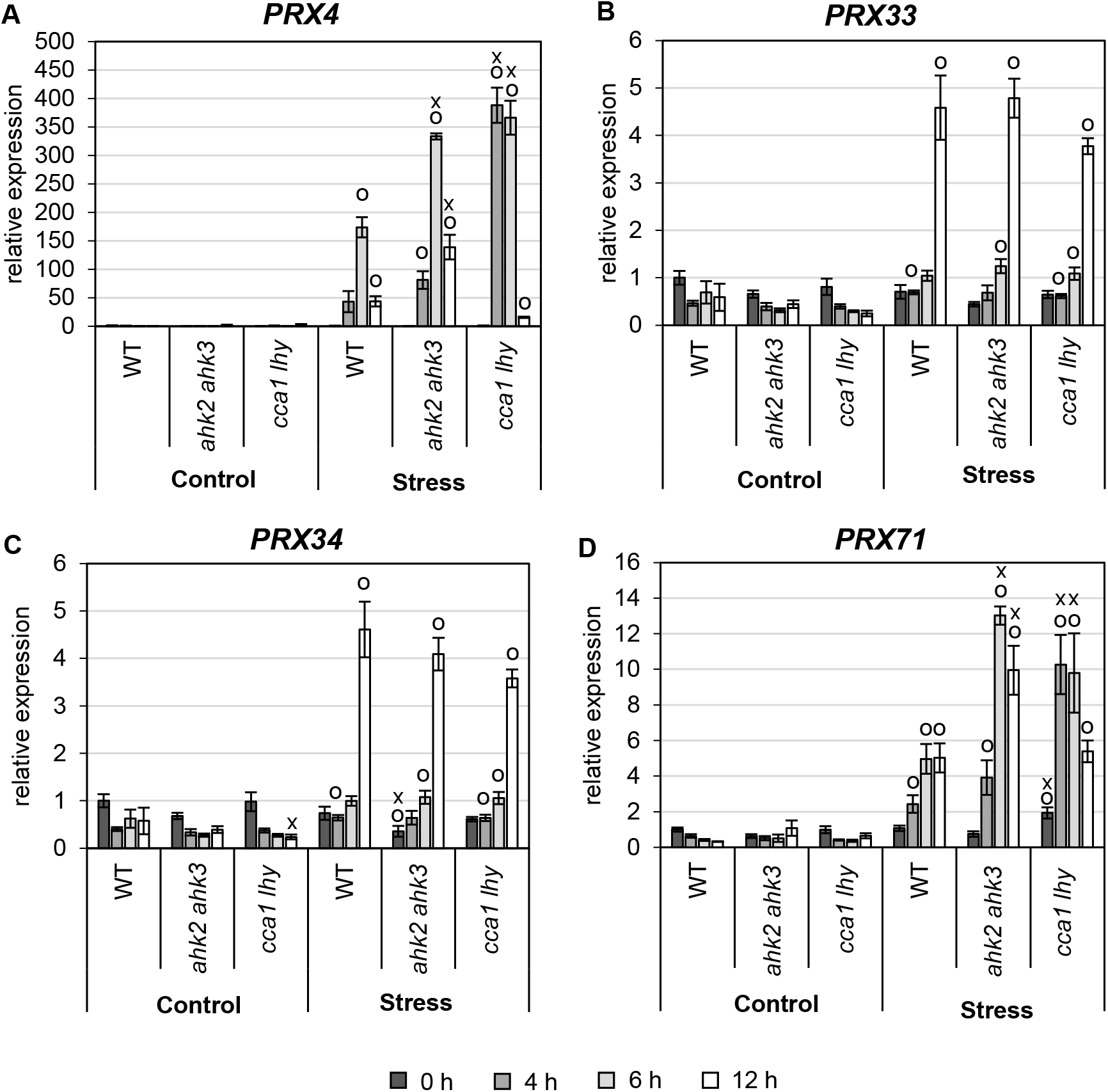
Expression of apoplastic peroxidase genes in response to photoperiod stress. Transcript abundances of *PEROXIDASE4* (*PRX4*) (A), *PRX33* (B), *PRX34* (C) and *PRX71* (D) in leaves under control conditions and in response to photoperiod stress at time points indicated in Fig. 6A. Expression levels were normalized to 0 h wild-type control, which was set to 1. Data are mean values ± SE (n = 4). Symbols indicate significant differences from the corresponding control (o) or the respective wild type under the same conditions and time point (x) (p-values < 0.05; *t-*test).

In addition to these genes encoding scavenging enzymes, the expression of genes coding key enzymes in the biosynthesis of the non-enzymatic antioxidants ascorbate, tocopherol and glutathione – namely the *VTC2*, *VTE1* and *GSH2* genes – were analyzed (Fig. 8). Under control conditions all genes showed a similar expression profile in all genotypes, with generally higher expression levels of *VTC2* and *VTE1* genes in *cca1 lhy*. Photoperiod stress treatment caused lowered transcript levels of the *VTC2* and *VTE1* genes as compared to control conditions. Suppression of the typical night elevation of the *VTC2* and *VTE1* gene expression might contribute to reduced levels of AsA, and eventually also of tocopherol, induced by photoperiod stress.

**Fig. 8.**
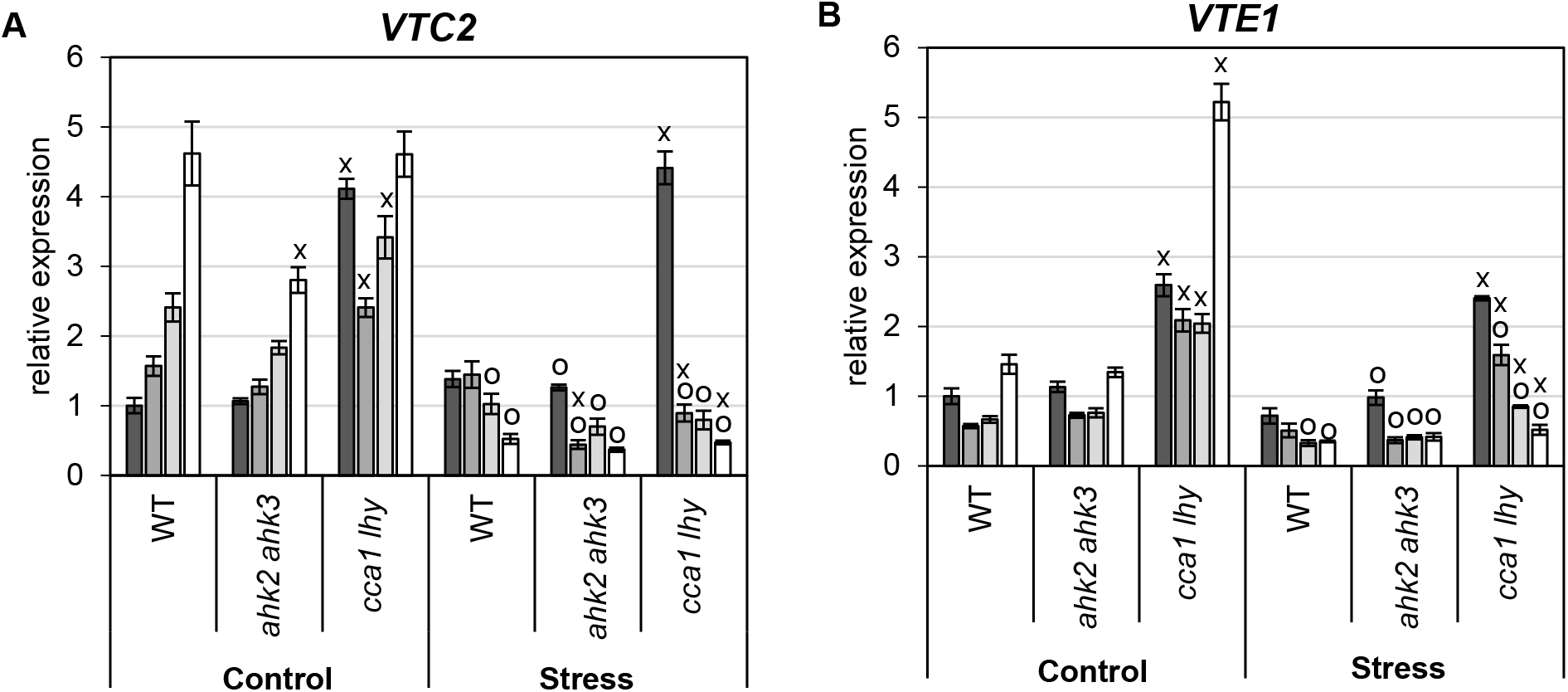
Regulation of transcripts of key genes in non-enzymatic antioxidants biosynthesis in response to photoperiod stress. Transcript abundance of *VTC* (A) and *VTE* (B) in leaves under control conditions and in response to photoperiod stress at time points indicated in Fig. 6A. Expression levels were normalized to 0 h wild-type control, which was set to 1. Data are mean values ± SE (n = 4). Symbols indicate significant differences from the corresponding control (o) or the respective wild type under the same conditions and time point (x) (p-values < 0.05; *t-*test).

## Discussion

This study has revealed several distinct changes of the cellular redox system to photoperiod stress. The photoperiod stress response was accompanied by a strong decrease in AsA redox (Fig. 1D) and the nightly increase in transcript levels of the AsA synthesis gene *VTC2* (Fig. 8A) was completely lacking after a photoperiod stress treatment. This suggested that a lowered AsA synthesis and therefore a lowered ROS buffering capacity might be part of the cause for the photoperiod stress syndrome. Noteworthy, AsA is found in the apoplast where it is the major non-enzymatic antioxidant (Shigeoka and Maruta, 2014). A reduction of the AsA content in this compartment would decrease the anyhow low antioxidant-buffering capacity of the apoplast even further (Podgorska *et al.*, 2017). The activities of the enzymes of the AsA-GSH scavenging system (APX, MDHAR, DHAR, GR) were not affected by photoperiod stress (Fig. 3, S2) indicating that the AsA-GSH cycle has no strong role in the stress response. The concentrations of phenolics and flavonoids were rather weakly altered by photoperiod stress (Fig. 2) like the no or only slight changes in SOD enzyme activity (Fig. 3; Suppl. Fig. S2A, B) excluding these cell internal components from being causative for the detrimental consequences of photoperiod stress.

### Oxidative stress by prolongation of the light period is associated with altered catalase and peroxidase activity

In contrast to the – with the exception of the AsA redox – rather minor changes in the non-enzymatic scavenging compunds, much stronger changes were noted in enzyme activities and transcript levels of genes involved in H_2_O_2_ metabolism and generation of peroxides (Fig. 4–7).

Changes in the activity of several enzymes might be a main cause for the oxidative stress. Catalase activity was rapidly reduced in all genotypes after beginning of the night to only about 20% of the original activity and remained low in the stress-sensitive genotypes (Fig. 4B) indicating a reduced capacity to detoxify H_2_O_2_.

Also PRXs showed an altered behavior in response to photoperiod stress. Total and apoplastic PRX activity increased at the middle of the night following an extended photoperiod The increase was consistent during the whole night and the following day in stress-sensitive genotypes. Consistent with the increase in enzyme activity, the expression of all four tested *PRX* genes was induced by photoperiod stress although with different response profiles. The fastest and strongest responses were shown by *PRX4* and *PRX71* with an enhanced induction in the *ahk2 ahk3* and *cca1 lhy* mutants (Fig. 7A, D). Their stronger induction might contribute to the stronger phenotypic consequences of photoperiod stress in these mutants. Additional *PRX* genes that are responsive to photoperiod stress and controlled by CK and/or the circadian clock are to be expected among the 73 class III peroxidase genes of *Arabidopsis* (Valerio *et al*., 2004). The strong induction of *PRX* genes and increase in PRX activity in response to photoperiod stress is a key finding contributiog to explain the destructive consequences of strong photoperiod stress.

### Cytokinin and the circadian clock are required to counteract the oxidative stress caused by photoperiod stress

Photoperiod stress clearly occurs in WT but the response to it was much stronger in CK-deficient plants (Nitschke *et al.*, 2016; this work). The stronger downregulation of catalase activity and the stronger induction of PRX activity in the *ahk2 ahk3* CK receptor mutant suggested a negative regulatory role of CK on the generation of oxidative stress. This is consistent with reports in the literature (reviewed by Cortleven *et al.*, 2019). For example, CK negatively regulates the formation of ROS in response to high light stress (Cortleven *et al.*, 2014) and crosstalk between ROS and CK is relevant to ensure proper functioning and maintenance of meristems in response to stress (Tognetti *et al.*, 2017). However, only little is known about the signaling pathways linking CK and oxidative stress. Genes encoding ROS scavenging proteins are among the most stably and rapidly CK-regulated genes suggesting that they could be direct targets of transcription factors mediating changes in CK signaling (Brenner and Schmülling, 2012, 2015). One direct link to regulate ROS formation by CK is through ARR2, a CK-regulated transcription factor known to bind directly to the promoters of the *PRX33* and *PRX34* genes (Arnaud *et al.*, 2017). Another CK-regulated transcription factor, CYTOKININ RESPONSE FACTOR6, is responsive to oxidative stress (Zwack *et al.*, 2013) and regulates crosstalk between H_2_O_2_ and the CK system (Zwack *et al.*, 2016). Notably, CK has anti-oxidative stress activity even in bacterial (Wang *et al.*, 2017) and human (Othman *et al.*, 2016) cells which suggests that the hormone might have acquired this function very early during evolution and retained it ever since in different organisms (Kabbara *et al.*, 2018).

Further, the data underpin that a functional circadian clock is required for a proper response to photoperiod stress. A tight link between the clock and oxidative stress is known (Lai *et al.*, 2012) and it has been proposed that clocks originally evolved to anticipate the presence of ROS (Edgar *et al.*, 2012). It has been suggested that CCA1 is a master regulator of ROS homeostasis through association with the Evening Element in promoters of ROS genes (Lai *et al.*, 2012). Loss of *CCA1* would lead to disturbance of the fine-tuned responses to oxidative stress and thus hamper the plants’ ability to properly master oxidative stress responses. Therefore, it is conceivable that the strong phenotypic consequences caused by photoperiod stress in the *cca1 lhy* mutant is due to improper clock output and altered transcriptional regulation of ROS genes.

## Conclusions

Together, this study shows that *Arabidopsis* has a response system to react to changes of the photoperiod. Although the experimental conditions do not occur in nature, we hypothesize that the nightly change in cellular redox status in response to photoperiod stress contributes to fine-tuning of plant responses to their environment. Naturally occurring changes in the day length due to seasonal shifts are in the range of few minutes per day, which might be too short to trigger any stress response. However, due to weather conditions or conditions of the habitat, plants perceive light of different quality and quantity throughout the day for longer time periods. Further, plants may be exposed to artificial light sources (e.g. street lights) causing extended photoperiods. Notably, an altered photoperiod might not necessarily cause harmful stress but a low stress level might also be beneficial since ROS are no longer seen solely as damaging side-products due to life in an O_2_-rich atmosphere but are also part of the cellular communication in plants with multiple beneficial functions (Mittler, 2017; Krasensky-Wrzaczek and Kangasjarvi, 2018; Noctor *et al.*, 2018).

## Supporting information

Abuelsoud et al_Supplemental data

## Supplementary information

**Supplemental Fig. S1.** Photoperiod stress response in leaves of WT, *ahk2 ahk3* and *cca1 lhy plants*.

**Supplemental Fig. S2.** Changes in enzymatic antioxidant activity in leaves in response to photoperiod stress.

**Supplemental Fig. S3.** Changes in activity of cell wall-bound peroxidase in leaves in response to photoperiod stress.

**Supplemental Table S1.** Sequences of primers used in this study.

**Supplemental Table S2.** Changes in concentrations of reduced AsA and total AsA in response to photoperiod stress.

**Supplemental Table S3.** Activity of glucose-6-phosphate dehydrogenase (G6PDH) in the apoplastic fluid.

## Acknowledgements

We thank Silvia Nitschke for critical review of the manuscript and acknowledge funding by Deutsche Forschungsgemeinschaft in the frame of Collaborative Research Centre 973 (www.sfb.973) and by grant Schm 814/27-1.

