## Supplementary material for "Photoperiod stress alters the cellular redox status and is associated with an increased peroxidase and decreased catalase activity": Abuelsoud et al_Supplemental data

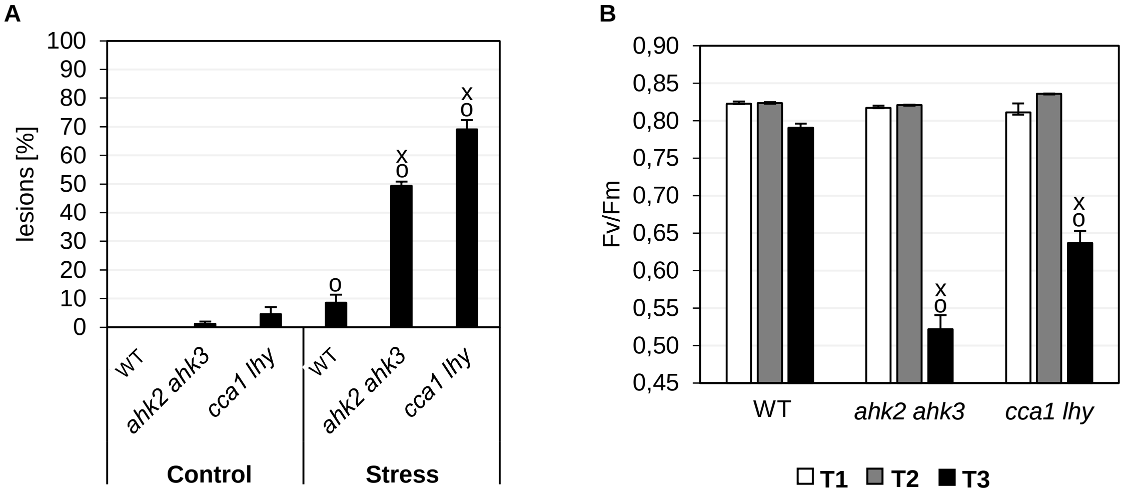


**Fig. S1.** Photoperiod stress response in leaves of WT, *ahk2 ahk3* and *cca1 lhy* plants. A schematic overview of the experimental setup is shown in Fig. 1A. (A) Percentage of leaves with lesions (n = 15) and (B) PSII maximum quantum efficiency (Fv/Fm) (n = 10). Data are mean values ± SE. Symbols indicate significant differences from corresponding control or plants at time point T1 (o) and the respective wild type under the same conditions and time point (x) (p-values < 0.05; *t*-test).

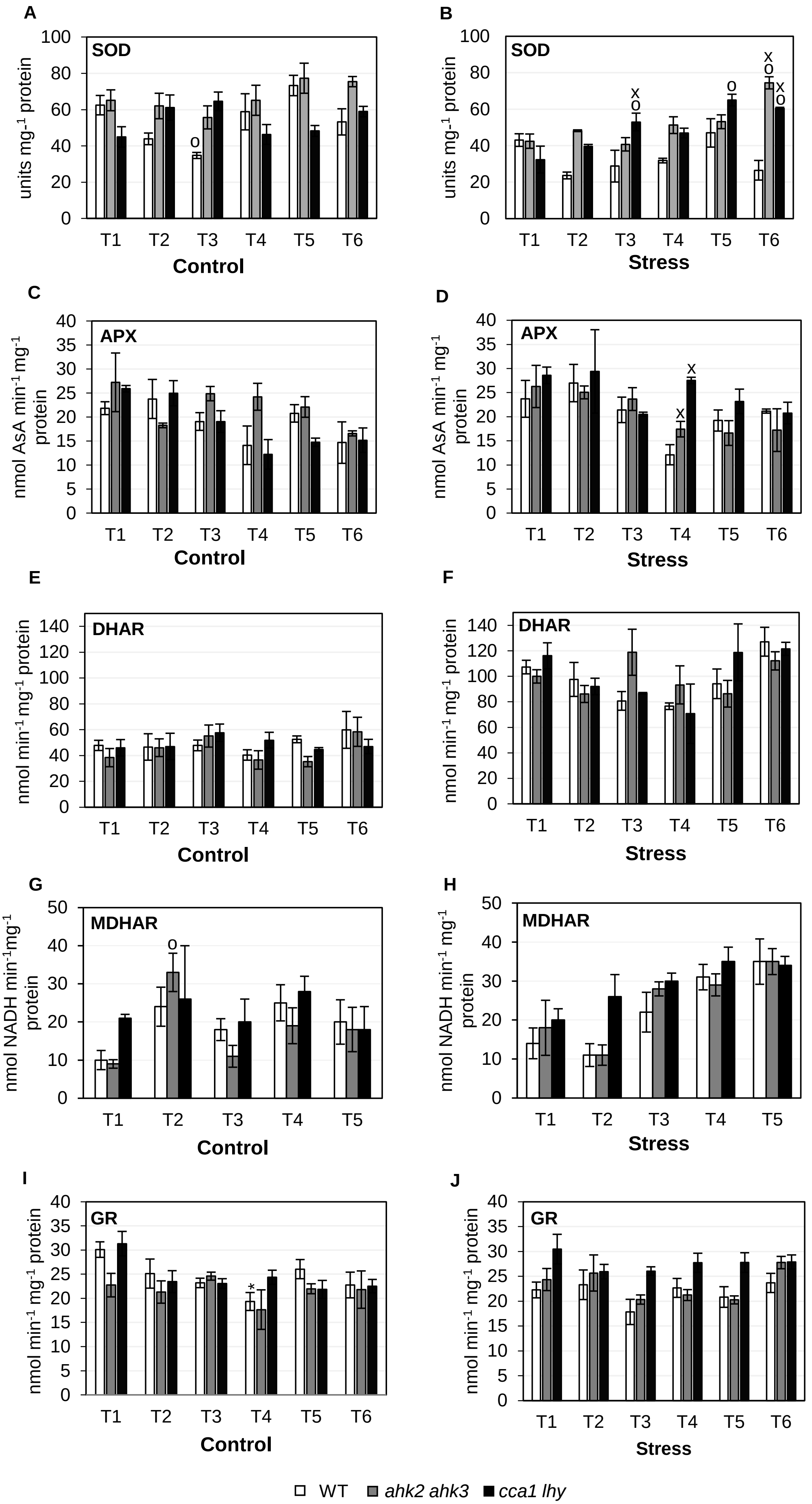

A
SOD
units mg-1 protein
T1 T2 T3 T4 T5 T6
Control
B
SOD
Stress
C
APX
nmol AsA min-1 mg-1 protein
D
APX
E
DHAR
nmol min-1 mg-1 protein
F
DHAR
G
MDHAR
nmol NADH min-1mg-1 protein
H
MDHAR
I
GR
J
GR
WT
ahk2 ahk3
cca1 lhy


**Fig. S2.**

**Fig. S2.** Changes in enzymatic antioxidant activity in leaves in response to photoperiod stress. Samples were collected during the night following a 32 h prolonged light treatment. Superoxide dismutase (SOD) (A, B), ascorbate peroxidase (APX) (C, D), dehydroascorbate reductase (DHAR) (E, F), monodehydroascorbate reductase (MDHAR) (G, H) and glutathione reductase (GR) (I, J) activities under control and stress conditions. Data are mean values (n = 4 ± SE). Symbols indicate significant differences from the corresponding plants at time point T1 (o) or the respective wild type under the same condition and time point (x) (p-values < 0.05; *t*-test).

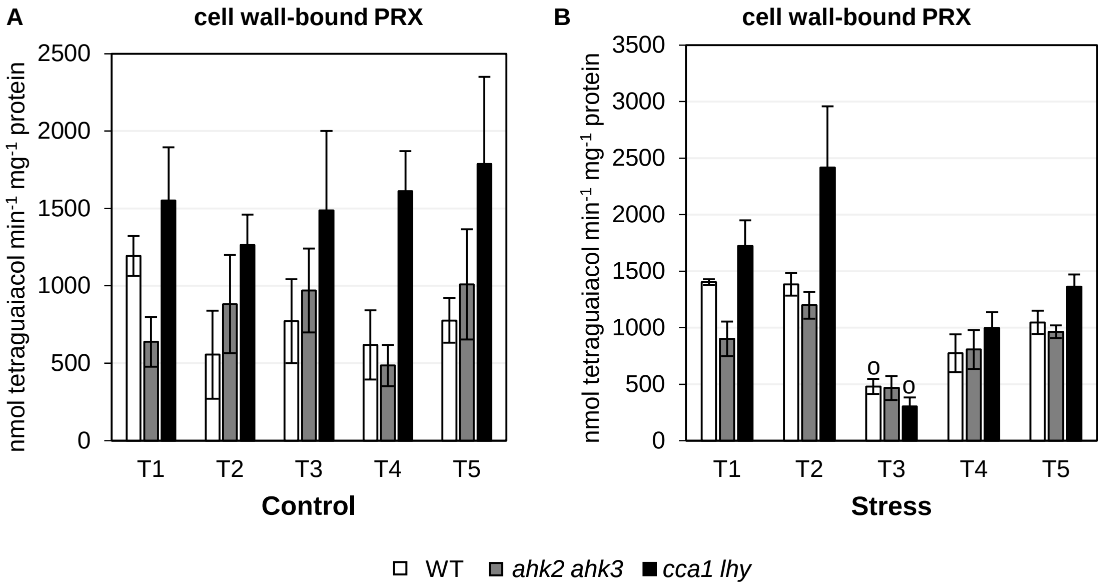


**Fig. S3.** Changes in activity of cell wall-bound peroxidase in leaves in response to photoperiod stress. Samples were collected at time points indicated in Fig. 4A under control (A) and stress (B) conditions. Symbols indicate significant differences from the corresponding plants at time point T1 (o) or the respective wild type under the same condition and time point (x) (p-values < 0.05; *t*-test).

**Table S1.** Sequences of primers used in this study.

| Gene name | ATG number | Forward primer | Reverse primer |
|---|---|---|---|
| *BAP1* | AT3G61190 | CCAGAGATTACGGCGCGTGTT | TACAGACCCCAAACCGGAACTCC |
| *CAT1* | AT1G20630 | AGGAGCCAATCACAGCC | TCAAGACCAAGCGACCA |
| *CAT2* | AT4G35090 | AACTCCGCCTGCTGTCTG | ATAGGGCATCAATCCATC |
| *CAT3* | AT1G20620 | TCACAGCCACGCCACTAA | AGAACCAAGCGACCAACC |
| *GSH2* | AT5G27380 | AGCAGTCGCAGTGGTTTATTTC | CCTCTTTGTTGTCCAGAAACCTC |
| *JAZ1* | AT1G19180 | CCCAACACCATTGACAGAAC | CTAAACCGAGCCACGACA |
| *LOX3* | AT1G17420 | ACGTTGTCGTACTGGTCGCC | GTCTCGTGGCACATACATAGGTAATG |
| *MCP2D* | AT1G79340 | AACCCGCTATGCAGACACACG | CAGTTGGTTTCCCCGCTGGA |
| *PP2A* | AT3G25800 | CCATTAGATCTTGTCTCTCTGCT | GACAAAACCCGTACCGAG |
| *PRX33* | AT3G49110 | AATCTGTCACTTTGGCAGGAG | GAATGGAGCTGGAAGATTTGCG |
| *PRX34* | AT3G49120 | TAGGGTCGGGTAAACCTGTG | GTTGCTCTCTCCGGTGGTC |
| *PRX4* | AT1G14540 | GGGCTAGCCTCAACGATCTC | AATCCCGCGTCAATGTCACT |
| *PRX71* | AT5G64120 | TCGTGACACAGTCATTCTCACTC | TTTCTGTTGTTGAACGGCAACG |
| *UBC21* | AT5G25760 | ACTCTTAGCCAAGTAGTGCTCC | GAATCACGGCCAACAATC |
| *VTC2* | AT4G26850 | GTTTCAGACTGCTGTGTTTGCC | GTTTCTCTGCGTAACACTGTGG |
| *VTE1* | AT4G32770 | ATTGCCTGGATTGACTGAGACC | GTTCTTGCCTCTAGTTCCACCAC |
| *ZAT12* | AT5G59820 | CGCTTTGTCGTCTGGATTG | AGCAGCCCCACTCTCGTT |

**Table S2.** Changes in levels of reduced AsA and total AsA in response to photoperiod stress. The experimental setup is shown in Fig. 1A. Data are mean values ± SE (n = 3). Symbols indicate significant differences from the corresponding control (o) or the respective wild type under the same conditions (x) (p-values < 0.05; *t*-test). FW, fresh weight. T1 corresponds to the beginning, T2 to the middle and T3 to the end of the night following photoperiod stress

| | | **Reduced AsA (μg g$^{-1}$ FW)** | | | **Total AsA (μg g$^{-1}$ FW)** | | |
|---|---|---|---|---|---|---|---|
| | | T1 | T2 | T3 | T1 | T2 | T3 |
| **Control** | WT | 321.4 ±6.9 | 352.8 ±23.7 | 370.9 ±8.4 | 339.0 ±7.7 | 468.5 ±19.1$^{o}$ | 466.1 ±20.0$^{o}$ |
| | *ahk2 ahk3* | 232.7 ±7.2 | 240.0 ±34.7 | 230.4 ±18.5 | 252.9 ±9.5 | 326.0 ±32.2 | 299.4 ±10.7 |
| | *cca1 lhy* | 266.7 ±20.2 | 232.4 ±12.7 | 293.3 ±28.8 | 283.3 ±19.1 | 325.3 ±9.7 | 367.7 ±32.7 |
| **Stress** | WT | 321.4 ±6.9 | 484.2 ±31.3* | 117.2 ±48.6$^{o*}$ | 339.0 ±7.7 | 635.5 ±27.7$^{o*}$ | 378.1 ±4.5* |
| | *ahk2 ahk3* | 232.7 ±7.2 | 375.8 ±22.6$^{o*}$ | 4.5 ±2.5$^{o*}$ | 252.9 ±9.5 | 466.5 ±27.3$^{o*}$ | 845.8 ±44.3$^{o*}$ |
| | *cca1 lhy* | 266.7 ±20.2 | 490.0 ±71.1$^{o*}$ | 54.8 ±2.4$^{o*}$ | 283.3 ±19.1 | 637.1 ±21.3$^{o*}$ | 551.2 ±73.3$^{o}$ |

**Table S3.** Activity of glucose-6-phosphate dehydrogenase (G6PDH) in the apoplastic fluid. Enzymatic activity is given as nmol $min^{-1}$ $mg^{-1}$ protein). Data are mean values of ± SE (n = 3). The experimental setup is shown in Fig. 6A.

| | **Control** | | | **Stress** | | |
|---|---|---|---|---|---|---|
| | WT | *ahk2 ahk3* | *cca1 lhy* | WT | *ahk2 ahk3* | *cca1 lhy* |
| **T1** | 0.10 ± 0.09 | 0.02 ± 0.01 | 0.04 ± 0.03 | 0.07 ± 0.06 | 0.00 | 0.06 ± 0.05 |
| **T2** | 0.00 | 0.00 | 0.04 ± 0.03 | 0.00 | 0.00 | 0.00 |
| **T3** | 0.03 ± 0.01 | 0.21 ± 0.17 | 0.34 ± 0.11 | 0.08 | 0.06 ± 0.05 | 0.00 |
| **T4** | 0.03 ± 0.02 | 0.12 ± 0.1 | 0.00 | 0.03 ± 0.02 | 0.07 ± 0.06 | 0.00 |
| **T5** | 0.03 ± 0.02 | 0.13 ± 0.04 | 0.08 | 0.32 ± 0.26 | 0.49 ± 0.31 | 0.09 ± 0.0 |
